# Genome-Wide Identification of Stable RNA Secondary Structures Across Multiple Organisms Using Chemical Probing Data: Insights into Short Structural Motifs and RNA-Targeting Therapeutics

**DOI:** 10.1101/2024.10.08.617329

**Authors:** Jingxin Wang

## Abstract

Small molecules targeting specific RNA binding sites, including stable and transient RNA structures, are emerging as effective pharmacological approaches for modulating gene expression. However, little is understood about how stable RNA secondary structures are shared across organisms, an important factor in controlling drug selectivity. In this study, I provide an analytical pipeline named RNA Secondary Structure Finder (R2S-Finder) to discover short, stable RNA structural motifs for humans, *Escherichia coli* (*E. coli*), SARS-CoV-2, and Zika virus by leveraging existing in vivo and in vitro genome-wide chemical RNA-probing datasets. I found several common features across organisms. For example, apart from the well-documented tetraloops, AU-rich tetraloops are widely present in different organisms. I also found that the 5’ untranslated region (UTR) contains a higher proportion of stable structures than the coding sequences in humans, SARS-CoV-2, and Zika virus. In general, stable structures predicted from in vitro (protein-free) and in vivo datasets are consistent in humans, *E. coli*, and SARS-CoV-2, indicating that most stable structure formation were driven by RNA folding alone, while a larger variation was found between in vitro and in vivo data with certain RNA types, such as human long intergenic non-coding RNAs (lincRNAs). Finally, I predicted stable three- and four-way RNA junctions that exist both in vivo and in vitro conditions, which can potentially serve as drug targets. All results of stable sequences, stem-loops, internal loops, bulges, and three- and four-way junctions have been collated in the R2S-Finder database (https://github.com/JingxinWangLab/R2S-Finder), which is coded in hyperlinked HTML pages and tabulated in CSV files.

## INTRODUCTION

Using small molecules to target RNA structures is an emerging pharmacological approach to regulate gene expression (1–4). Unlike antisense oligonucleotides, which recognize the primary sequence by Watson–Crick base-pairing, small molecules recognize various binding pockets formed during RNA folding. Although molecules have been developed to target tertiary structures (e.g., pseudoknots (5, 6), G-quadruplexes (7), and riboswitches (8)), dynamic structures in conformational equilibrium (9, 10), or transiently formed structures (11–14), many small molecules target short, stable RNA secondary structures, such as stem-loops, internal loops, and bulges (15–19). Characterizing these stable structural elements in different organisms is crucial to understand the selectivity and off-target effects of therapeutic agents targeting RNA secondary structures.

The current gold standard for probing RNA structures (in vivo and in vitro) at a scale of hundreds of nucleotides or more is chemical probing methods using 2’-OH acylation (e.g., selective 2’-hydroxyl acylation analyzed by primer extension [SHAPE]) (20–22) or nucleobase alkylation (e.g., methylation with dimethyl sulfate [DMS]) (23–26). Generally, SHAPE reagents and DMS are more likely to leave a covalent tag at unpaired RNA regions compared to paired regions, due to reaction accessibility, which can be quantified by stop or mutation during reverse transcription (20, 24, 27). As a result, low SHAPE and DMS scores can usually be correlated to high base-pairing probability, providing data from which secondary and higher-order RNA structures can be deduced. Advancements in chemical probing techniques, such as “clickable” groups on chemical probing reagents, enable enrichment of the chemically marked RNA, rendering a significant enhancement of probing signals for high-throughput studies (28). When coupled with next-generation sequencing, these advancements allow for RNA structure elucidation on a large scale, such as the whole human transcriptome (28, 29). Recently, almost all available genome-wide chemical probing data were collected in a single source, namely RNA Atlas of Structure Probing (RASP) (30). The increasing amount of chemical probing data has been utilized for various purposes. For example, SHAPEwarp was developed as a method to compare new SHAPE profiles with existing ones to study RNA structure conservation and similarities (31). In addition, several databases of RNA mapping across different organisms exist (32–34). However, most of these comprehensive databases are focused on RNA mapping data curation, processing, and application, but not categorizing defined structures. While the main purpose of the database reported in this paper is to compare structures across organisms, a database using structure-based categorization has the additional advantage of eliminating variance across chemical probing conditions (e.g., DMS vs SHAPE) and batches, facilitating comparisons across experiments.

By leveraging chemical probing data from RASP, I created a structure identification pipeline named RNA Secondary Structure Finder (R2S-Finder) that analyzes SHAPE scores to identify and characterize stable RNA structures such as RNA stem-loops, internal loops, bulges, and three- and four-way junctions. Here, I present a new database as the outcome of R2S-Finder, which contains the stable RNA secondary structures from published genome-wide chemical probing sequencing datasets for several well-studied organisms, including human (35), *Escherichia coli* (*E. coli*) (26), SARS-CoV-2 (36), and Zika virus (37). The chosen organisms may or may not represent other similar species (e.g., other viruses), but have been commonly studied and provide proof-of-principle for comparing structures across organisms. On top of the established method using low SHAPE/low entropy metrics to determine the stable regions (38), my analysis had stringent criteria limiting alternative structures within the stable region (Figure 1), making the structure classification unambiguous (see Material and Methods). The resulting R2S-Finder database is coded in HyperText Markup Language (HTML) and tabulated in comma-separated values (CSV) files, which allow users to browse organisms and genes or perform further analysis. The R2S-Finder database also allows users to search stable RNA secondary structures for potential drug targets and make preliminary predictions for drug selectivity by comparing structures across organisms. The R2S-Finder pipeline reported here can also be used to evaluate chemical probing data on other organisms or updated datasets.

**Figure 1.**
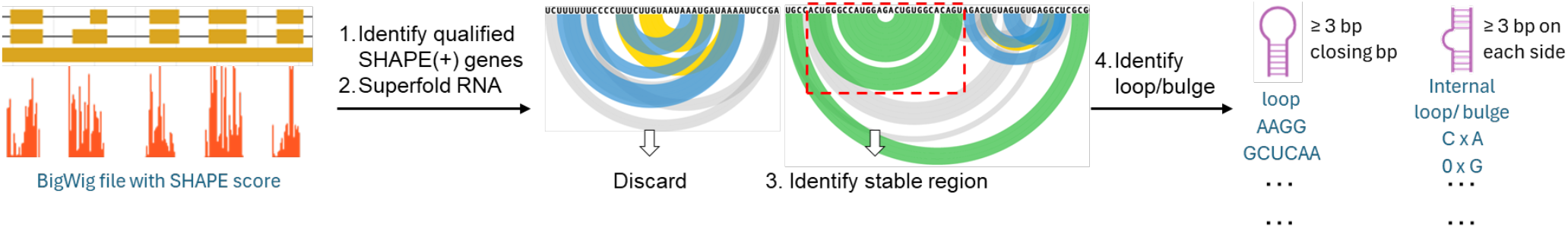
Stable RNA structure determination in R2S-Finder pipeline. The workflow contains four steps: (1) Identification of genes with qualified SHAPE score from the BigWig files. (2) Prediction of RNA structures and base-pairing probabilities using Superfold software package in batches. The colour of the arcs in the arc plot represents the base-pairing probabilities: green (80–100%), blue (30–80%), yellow (10–30%), and grey (<10%). (3) Identification of stable RNA structures from base-pairing probabilities. (4) Identification of stem-loop, internal loop, and bulge structures within the stable RNA sequences.

## MATERIAL AND METHODS

### R2S-Finder: Chemical probing signal extraction and stable RNA structure identification

The workflow for stable RNA structure identification is summarized in Figure 1. In this paper, most analyses used SHAPE data, except the *E. coli* rRNA analysis, which used DMS data with a nearly identical pipeline as I describe below. In step 1, I downloaded SHAPE scores associated with genome annotations from the RASP database for human, *E. coli*, SARS-CoV-2, and Zika virus. I extracted genes with valid SHAPE scores (including zero reactivity) of at least 11 nucleotides (nts) in length (denoted as SHAPE(+) regions). If the SHAPE(+) regions had 5% missing SHAPE signals within the region, I discarded the genes (or transcripts) without further analysis. For DMS data, the tolerability of missing data was increased from 5% to 65%, because G and U are considered DMS(–). I reformatted the filtered data into SHAPE map files compatible with the Superfold software package (version 1.0) (https://github.com/Weeks-UNC/Superfold) (22, 39). The virus genomes were considered as single entries without separating the genes. The total number of SHAPE(+) nucleotides included in genes, along with the number of genes (as applicable), stable sequences, loops, internal loops, bulges, and three- and four-way junctions, are summarized in Table 1 for each organism.

**Table 1.** Statistics of the data used in this study with valid SHAPE data (in vivo / in vitro data).

|  | SHAPE (+) nts | # of genes | # of stable sequences | # of stem-loops | # of internal loops/bulges | # of 3/4-way junctions |
| --- | --- | --- | --- | --- | --- | --- |
| Human | 713,398 / 708,080 | 1,313 / 1,247 | 5,774 / 5,995 | 5,909 / 6,209 | 3,485 / 4,080 | 136 / 164 |
| *E. coli* | 181,733 / 188,443 | 223 / 225 | 1,703 / 1,835 | 1,774 / 1,938 | 1,457 / 1,877 | 42 / 48 |
| SARS-CoV-2 | 29,871 / 29,873 | – | 383 / 353 | 402 / 375 | 364 / 310 | 17 / 18 |
| Zika virus | 10,759 / 10,759 | – | 86 / 93 | 90 / 94 | 62 / 74 | 3 / 1 |

In step 2, RNA folding prediction was performed with all SHAPE map files using Superfold, which set the SHAPE pseudoenergy restraints to match the observed SHAPE signals. Superfold calls the Fold program in the RNAstructure package to calculate the Shannon entropy. Superfold/Fold also provides the probability of all base pairs (dp file) and the most probable RNA secondary structures (ct file).

To identify stable RNA regions in step 3, I used three criteria to remove ambiguous or unstable structures: (1) The most probable base pairing of any nucleotide must be contained within the region. (2) All nucleotides involved in base-pairing must have a probability > 10% (−log(Probability) < 1) at all points and an average probability > 50% (−log(Probability) < 0.301) across the entire region. (3) If such a stable region is fully included within a larger region that also fulfills the above two criteria, the larger stable region is used.

As the 4^th^ step in the pipeline, I defined stem-loops as the unpaired sequences of 3−10 nts with one stem having a minimum of 3 base pairs (bp). Similarly, internal loops and bulges were defined as unpaired sequences (0−10 nts on either side) with two stems having a minimum of 3 base pairs each. The internal loops and bulges are denoted as m × n and m × 0 (or 0 × n), respectively, where m is the unpaired nucleotides at the 5’ end and n at the 3’ end. It is worth noting that m × n bulges were considered equivalent to n × m bulges (e.g., AG × G and G × AG bulges) in my analyses. I mapped the sequences with SHAPE scores, along with the identified stable sequences, stem-loops, internal loops, and bulges, to reference transcript sequences of the specific organisms. I provide the gene names, transcript and gene IDs, stable sequences, bulges, and loops in hyperlinked HTML webpages as well as spreadsheets (CSV files) (https://github.com/JingxinWangLab/R2S-Finder).

### Coding and non-coding sequence extraction

For human RNA transcripts, if more than one isoform was available and eligible for R2S-Finder analysis, I chose the longest annotated transcript (i.e., full-length transcripts). For human data (in vivo SHAPE), I extracted the coding sequence (CDS) and untranslated region (UTRs) for protein-coding genes according to the human genome annotation file (GFF3 file, available at https://www.ensembl.org/info/data/ftp/index.html). For SARS-CoV-2 and Zika virus data (in vivo SHAPE), the CDS and UTRs are annotated based on the reference sequences (RefSeq) with accession numbers NC_045512.2 and NC_012532.1, respectively. Data points without any stable sequences were discarded during analysis.

### Three- and four-way junction RNA identification

I defined three- and four-way junctions as stable sequences with 2 and 3 stem-loops, respectively. I further extracted the 2D structures of three- and four-way junctions from the in vivo and in vitro datasets and reported them in dot-bracket notation (available at https://github.com/JingxinWangLab/R2S-Finder/tree/main/CSV_files). I predicted the 3D structures using transform-restrained Rosetta for RNA (trRosettaRNA) algorithms (40). The potential drug binding sites were also predicted using the PDB files generated by trRosettaRNA and Fpocket package with default settings (41).

## RESULTS

### General features of stable RNA secondary structure datasets

The R2S-Finder pipeline pinpoints stable sequences using chemical probing data and an integrated RNA folding prediction tool, Superfold, which is tailored for chemical probing data analysis (22, 39) (Figure 1). Essentially, the RNA folding prediction in Superfold was performed based on the Fold program in the RNAstructure software package, which is widely used for predicting and analyzing RNA secondary structures using thermodynamic models compatible with chemical probing constraints (42). Importantly, to categorize RNA secondary structures, R2S-Finder used stringent criteria (see Material and Methods) and discarded regions with ambiguous base pairs or multiple RNA conformations in equilibrium (Figure 1). As a result, R2S-Finder is a robust approach to identify short RNA structure motifs. I first applied the R2S-Finder algorithm on a well-studied gene, 16S rRNA in *E. coli* using an in-cell DMS dataset (Figure 2A) (43). The base-pairing prediction is generally consistent with that observed with reported SHAPE data (44). It is expected that R2S-Finder excludes some stable base pairs due to ambiguous structures (Figure 2A). Next, I inferred the location and sequence of stable structures across humans, *E. coli*, SARS-CoV-2, and Zika viruses in vitro and in vivo using R2S-Finder. I further identified stem-loops, internal loops, bulges, and three- and four-way junction structures within these stable sequences (Table 1). I made all structure results, along with the corresponding transcript or gene reference sequences, freely available for download in the R2S-Finder database in HTML and CSV formats. It is worthwhile noting that in the current genome-wide datasets I chose to analyse, some RNA types are not well represented in the chemical probing studies due to the sample preparation methods, such as human ribosomal RNAs and microRNAs.

**Figure 2.**
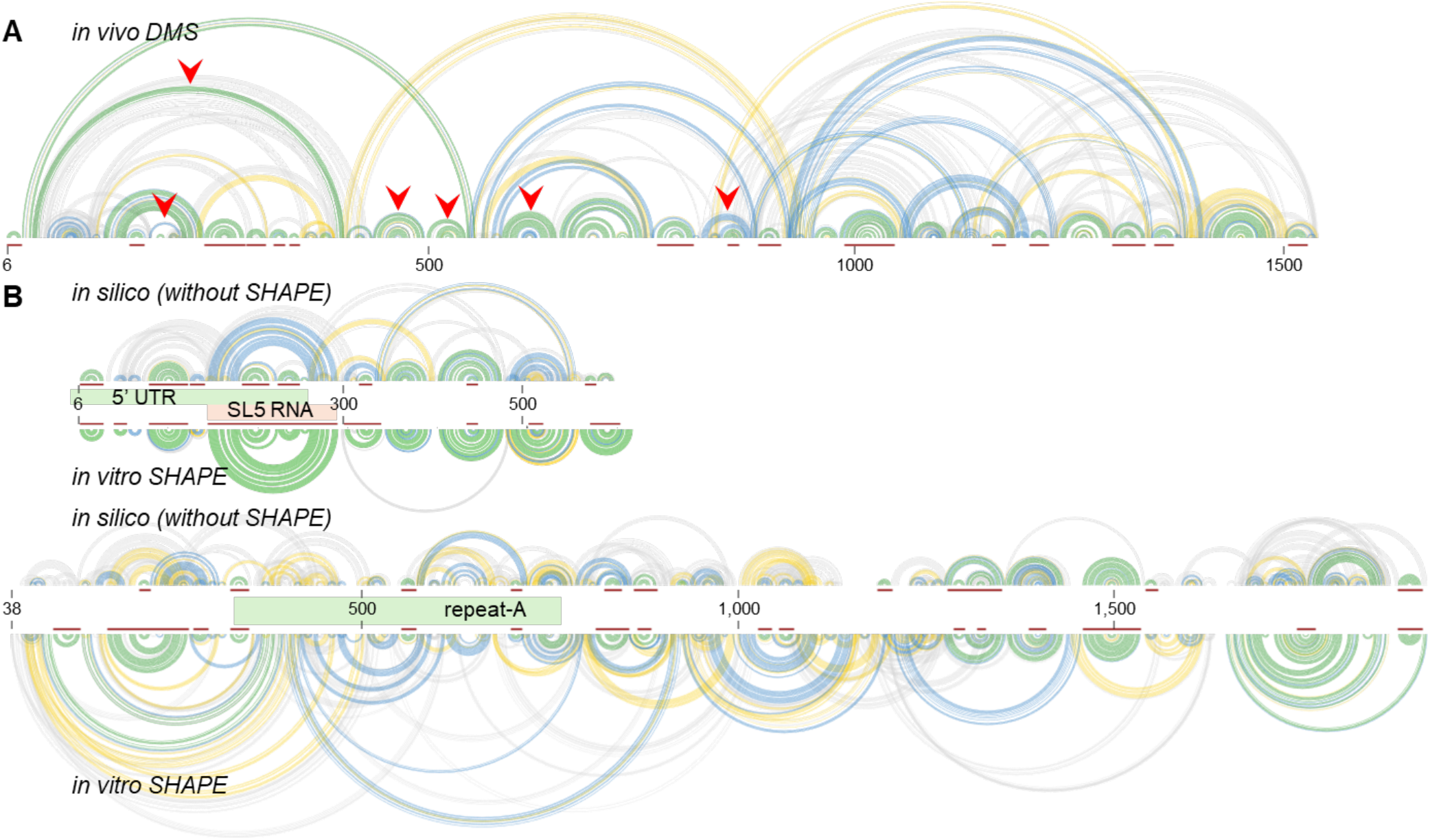
**(A)** Stable sequences identified by R2S-Finder for *E. coli* 16S rRNA (*E. coli* gene b0201). In the arc plot, the base-pairing probability is represented in green (80–100%), blue (30–80%), yellow (10–30%), or grey (<10%). The identified stable sequences are highlighted in red bars below the arcs. Red arrows indicate base pairs where ambiguous structures or multiple conformations in equilibrium exist within the base-paring range, which would not pass R2S-Finder’s selection criteria. **(B)** Comparison of RNA folding in the presence (in vitro) and absence (in silico) of SHAPE pseudoenergy restraints using Fold program with the 5’-end sequences in SARS-CoV-2 (top) and Xist (bottom) genes.

The Fold program uses pre-set thermodynamic parameters to determine the stability of the predicted structures that are usually accurate for short motifs, such as those analysed here. To investigate the effects of SHAPE pseudoenergy restraints on R2S-Finder results, I removed them for representative RNAs (i.e., highly stable SARS-CoV-2 gene and highly variable X-inactive specific transcript [Xist] gene, see discussion below) and evaluated changes in predicted stable sequences. In the stable region of the SARS-CoV-2 gene, the in silico RNA folding prediction was similar to that observed with SHAPE pseudoenergy restraints. However, the stable structure selection results using R2S-Finder were distinct using in vitro SHAPE data from in silico prediction due to the experiment-based base-pairing probability calculations (Figure 2B). The difference in R2S-Finder results between the in silico and in vitro SHAPE data is even more significant within the variable region of the Xist gene (Figure 2B). These results suggest that the SHAPE pseudoenergy restraints obtained in chemical probing experiments are crucial for short structural motif identification in R2S-Finder pipeline.

### Occurrence of stable structures, stem-loops, internal loops, and bulges

As discussed previously, due to the stringent criteria used to identify stable structures using chemical probing data (see Methods and Materials), short structural motifs were more likely to be selected by R2S-Finder. For all organisms in this study, most of the identified stable sequences are less than 30 nucleotides (nts) (Figure 3A). Long sequences (> 50 nts) comprise only 9–13% of identified stable sequences in humans, *E. coli*, SARS-CoV-2, and Zika virus. Next, I investigated the size of loops. Consistent with the literature, I found that tetraloops are more prevalent than other loop sizes across organisms, constituting 26–34% of all stem-loops with an occurrence rate of 2.6–3.9 per 1,000 nts (Figure 3B). In general, longer loops are less common. Finally, I found that the 1×0 bulges and 1×1 internal loops are significantly more common than other bulge or internal loop sizes, accounting for approximately 50% of all bulge/internal loop structures combined (Figure 3C). Notably, many small molecules optimized for drug development target small internal loops or bulges (45), highlighting the importance of selectivity in these contexts.

**Figure 3.**
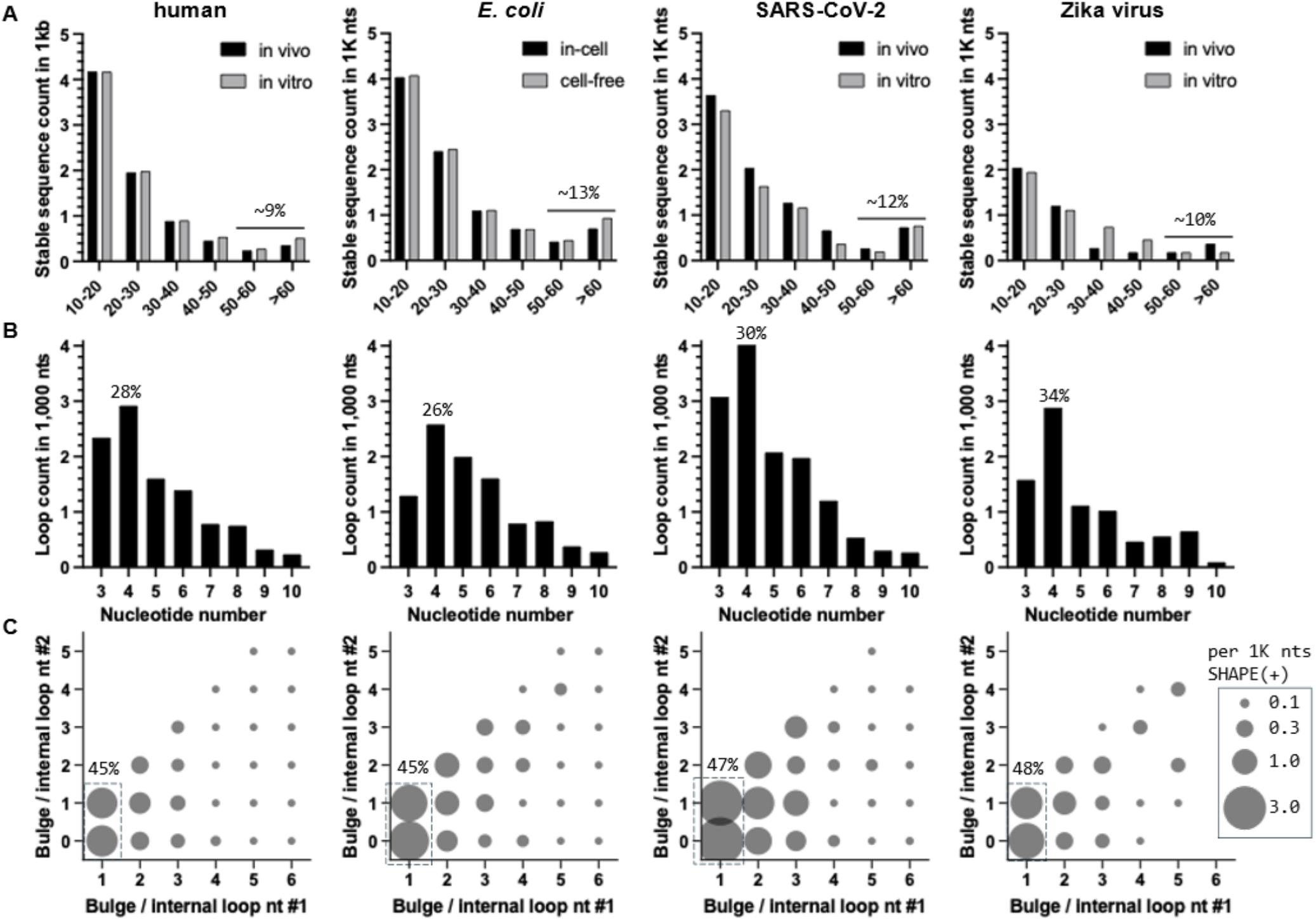
Length of stable sequences and sizes of loops and bulges in different organisms including humans, *E. coli*, SARS-CoV-2 and Zika virus. **(A)** Distribution of the length of the stable sequences. Both in vivo and in vitro data are included. The percentage of long sequences (> 50 nts) is annotated (average of in vivo and in vitro data). **(B)** Distribution of the number of loops with different sizes of the loop for in vivo data. The percentage of tetraloops is annotated. **(C)** Distribution of m × n internal loops and bulges for in vivo data, where m and n are the unpaired nucleotides in either strand of the stem. The diameter of the bubble represents the count in 1,000 SHAPE (+) nts. The percentage of 1×0 bulges and 1×1 internal loops combined is annotated.

I further analyzed various common tetraloop sequences reported in the literature, including GNRA (46), UNCG (47), CUYG (46), GANC (48), AGNN (49), and UYUN (50, 51), where N is any base, R is a purine, and Y is a pyrimidine. Some of these tetraloops were initially discovered in highly folded RNAs, such as bacterial rRNA and yeast Group II intron. I found that GNRA and AGNN are the two most prevalent tetraloops in the organisms studied (Figure 4A). Following these two tetraloops, I also frequently observed UYUN, identified as RNA-binding site for the histone mRNA stem-loop binding protein (50). While the majority of the tetraloops do not belong to any known classes, the top three unclassified tetraloops were all AU-rich—an unprecedented observation. The Zika virus was excluded from the analysis of the unclassified tetraloops due to a low total loop count. To verify the novel AU-rich tetraloops, I cross-checked the SHAPE scores in the loop regions and compared them to the 3 bp closing stems of the AU-rich tetraloops, where the SHAPE scores in AU-rich loops were significantly higher than in the stems, confirming their existence (Figures 4B, 4C). Interestingly, the closing base pairs of these tetraloops differ between humans and *E. coli*. It is well-recognized that GC base pair can provide favorable thermostability to tetraloops, especially GNRA and UNCG tetraloops (52). Indeed, in *E. coli*, GC base pairs are the most common closing base pairs among all the tetraloops tested. However, I observed that AU base pairs are more common in GNRA and AU-rich tetraloops in humans. This difference in closing base pairs between humans and *E. coli* may indicate that maintaining the stability of these RNA stem-loop structures is generally less essential in humans.

**Figure 4.**
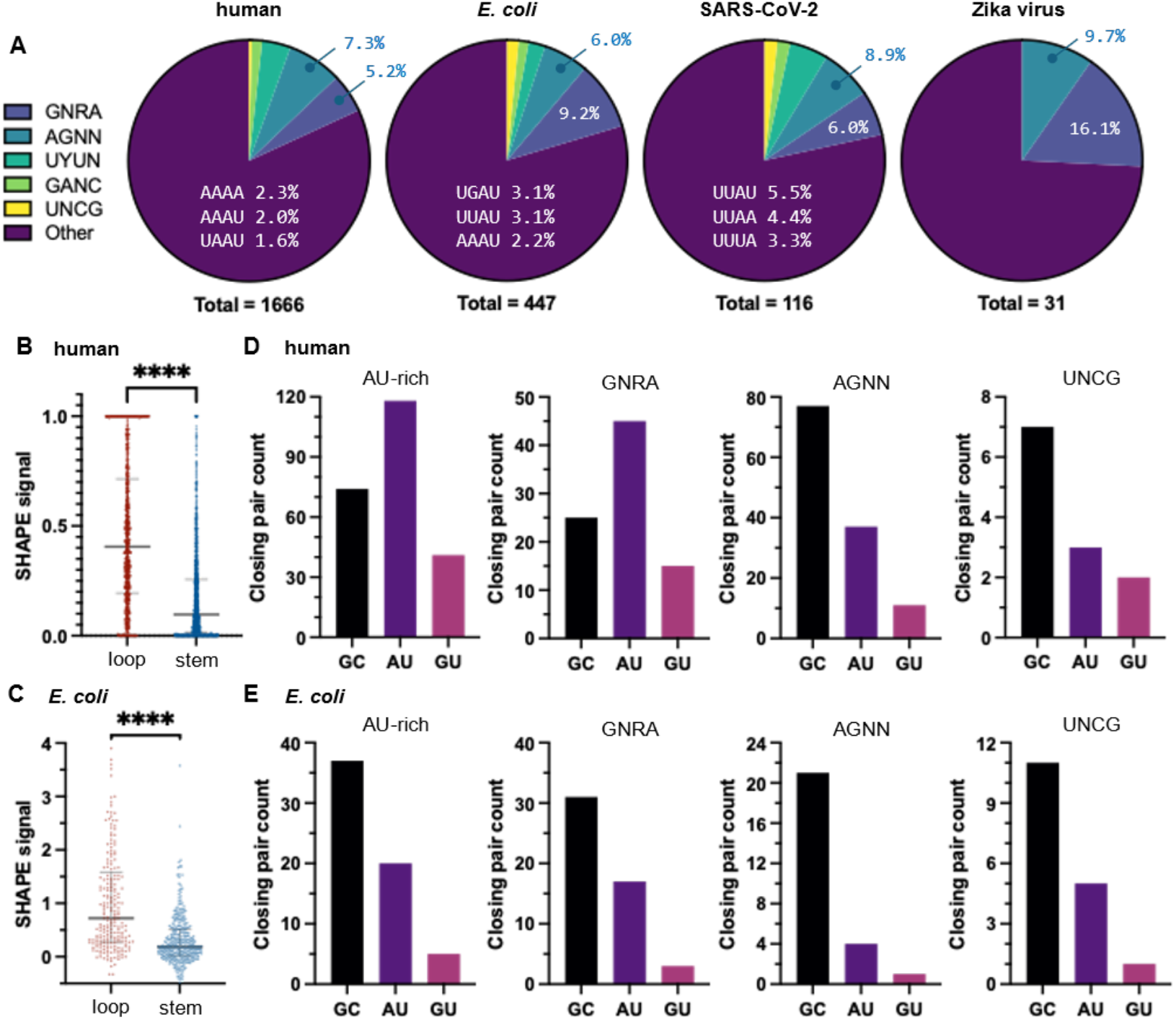
Prevalence of tetraloops and closing base pairs in humans, E. coli, SARS-CoV-2 and Zika virus using in vivo data. **(A)** Distribution of various tetraloops. The percentage of GNRA, AGNN, and top 3 unclassified tetraloops are annotated. Zika virus was excluded from the unclassified tetraloops analysis due to low total loop count. **(B–C)** SHAPE scores of the AU-rich tetraloops that only contain A and/or U and that of their 3 bp closing stems in (B) humans and (C) *E. coli* using in vivo data. **** indicated that p < 0.0001. **(D–E)** Analysis of the closing base pair of the common tetraloops in (D) humans and (E) *E. coli* using in vivo data.

Among 1×0 bulges, G×0 bulges are generally less prevalent than other types of bulges with the exception of SARS-CoV-2. In humans, G×0 bulges constitute only 14% of all 1×0 bulges, whereas A×0 bulges constitute ∼40% (Figure 5A). I observed a general ranking of bulge counts in the order G×0 < C×0 < U×0 < A×0 across most organisms in this study. For 1×1 internal loops, C×C are rarer than other types of 1×1 internal loops across organisms. In humans, C×C internal loops make up only 4.2% of 1×1 bulges, compared to the expected 10% if the distribution were random, and I found no C×C bulges in the stable sequences for Zika virus (Figure 5B). In small-molecule drug development, targeting secondary structures that are less common in humans, such as G×0 bulges or C×C internal loops, may reduce the likelihood of off-target effects, as these structures are less likely to be present in unintended RNA targets.

**Figure 5.**
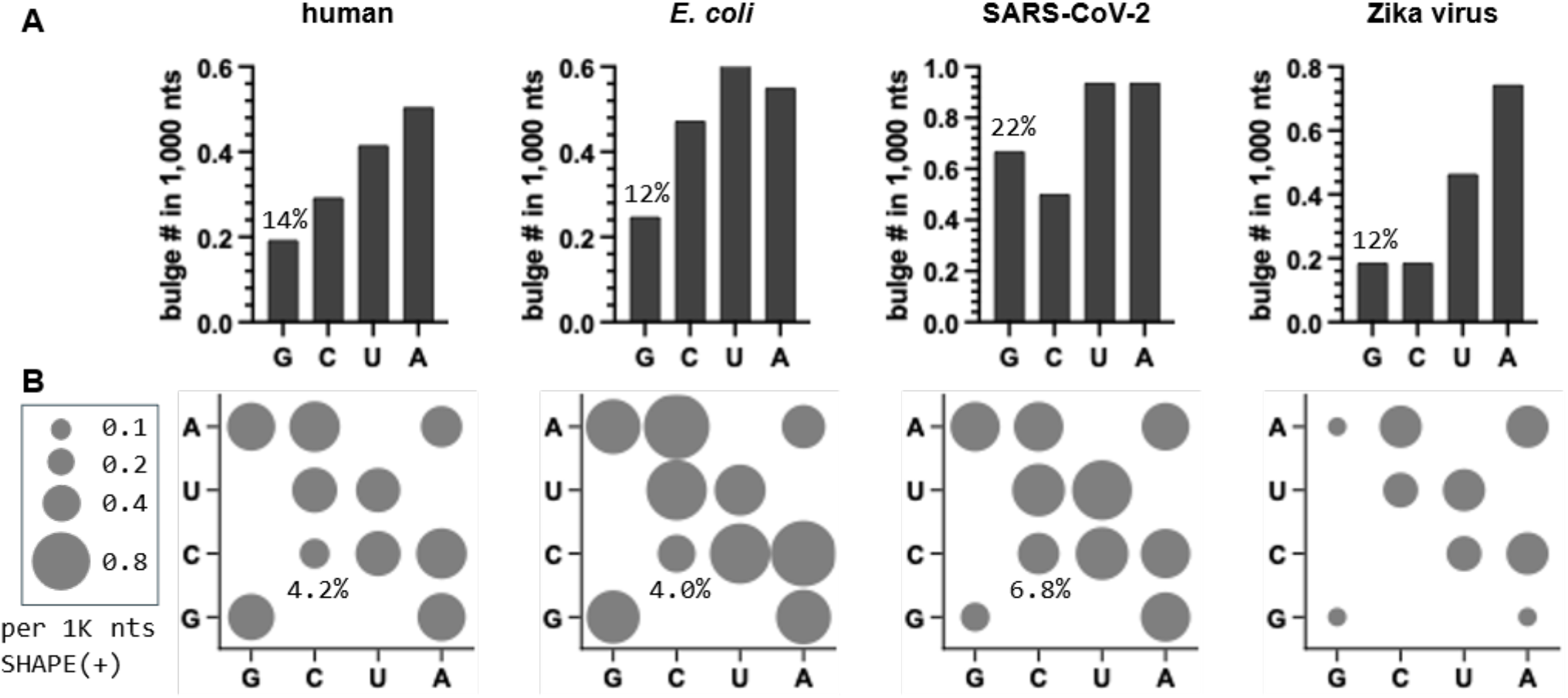
Prevalence of 1×0 bulges and 1×1 internal loops in different organisms including humans, *E. coli*, SARS-CoV-2, and Zika virus using in vivo data. **(A)** Distribution of 1×0 bulges. The percentage of G×0 tetraloops is annotated. **(B)** Distribution of 1×1 internal loops. The diameter of the bubble represents the count in 1,000 SHAPE(+) nts. The percentage of C×C bulge is annotated. A×U/U×A, C×G/G×C, G×U/U×G bulges do not exist due to Watson–Crick or wobble base pairing.

### Stable structure proportions comparing coding and non-coding regions

It has been previously demonstrated that 5’ untranslated regions (UTRs) in eukaryotic protein-coding genes use various RNA structures to regulate translation as these regions are involved in initiating protein synthesis (53). Therefore, I speculated that the 5’ UTRs of human and viral genes generally have more stable structures than the coding sequences (CDSs). I compared the lengths of stable sequences obtained via R2S-Finder in coding and non-coding regions. In humans, the 5’ UTR appears to have a higher proportion of stable sequences than CDS, while the stable sequence proportions within the CDS and 3’ UTR are similar (Figure 6A). Both SARS-CoV-2 and Zika virus are positive-sense single-stranded RNA (+ssRNA) viruses; their RNA genomes show that the 5’ UTR has more than twice the stable sequence ratio compared to the CDS (Figure 6B). In SARS-CoV-2, the 3’ UTR is considerably more flexible than the CDS region, where no short stable structural motif was identified by R2S-Finder, consistent with the large variable region which has different conformations in equilibrium (54). In Zika virus, the 3’ UTR is slightly more stable than the CDS, largely due to the presence of well-characterized stable structures, such as two large exoribonuclease-resistant RNA (xrRNA) elements (55), which protect the viral RNA from degradation— ensuring the stability of the viral genome during replication. However, the stable sequence ratio is still significantly lower than in the 5’ UTR (Figure 6B). This finding highlights the potential of targeting 5’ UTRs in viral genomes for antiviral therapies, as RNA stability in these regions may influence multiple aspects of virus life cycles (56–58).

**Figure 6.**
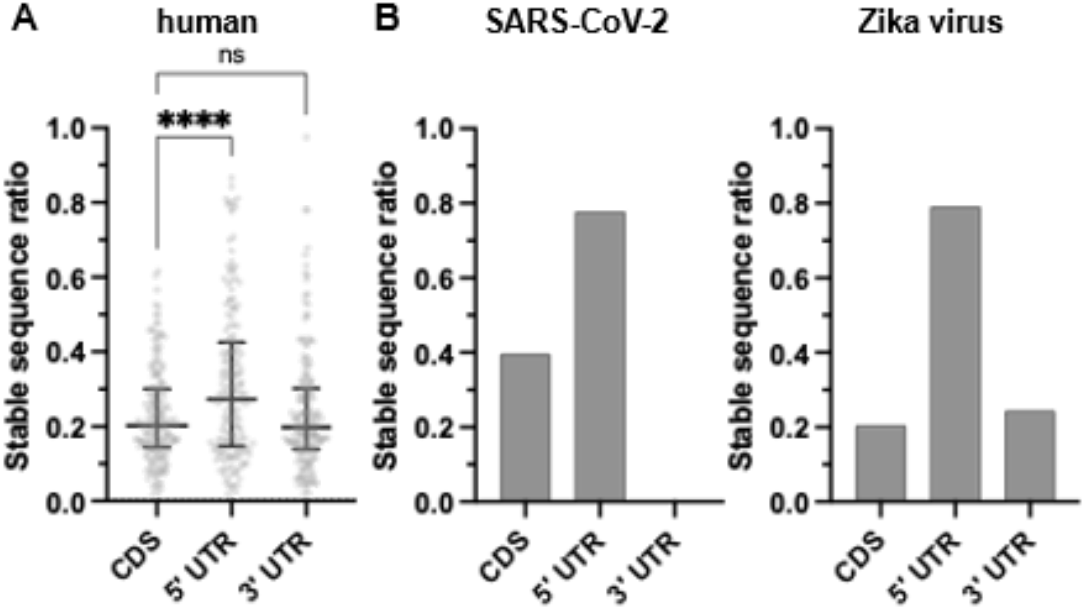
Stable sequences in coding and non-coding regions. Stable sequence ratio (stable sequence length / SHAPE(+) length in each category) in (A) humans and (B) viruses, including SARS-CoV-2 and Zika virus. In (A), the center horizontal line is the median of the data, and the error bar indicates the interquartile range. **** indicates p < 0.0001, ns = not significant.

### RNA secondary structure differences in vivo vs in vitro

It is well-known that in vivo and in vitro RNA structures differ (59) mainly due to the lack of RNA binding proteins in the protein-free in vitro experiments. Such differences captured by chemical probing experiments can be exploited to predict extensive protein-binding regions in RNA (59). I contend that the percentage of shared stable structures between in vivo and in vitro settings generally reflects the extent of RNA–binding protein interactions in cells. I examined the stable RNA secondary structures from the R2S-Finder pipeline and identified stable structures where predicted folding was retained in both in vivo and in vitro. I found that for both human and *E. coli* transcriptomes, ∼76% of the stable structures are shared between in vivo and in vitro (Figure 7A). The high percentage of shared structures suggests that these regions are primarily governed by RNA folding alone, rather than interactions with RNA-binding proteins or other cellular factors. Zika virus has a smaller number of shared stable sequences (67%), implying that Zika virus RNA might have more RNA–binding protein interactions in host cells. SARS-CoV-2 has the highest percentage of common structures (81%) in vivo and in vitro. In fact, in several hundreds of nucleotides at the 5’-end of the SARS-CoV-2 genome, the base-paring events under in vivo and in vitro conditions are almost identical (Figure 7B). Previously, we also compared chemical probing data in SARS-CoV-2 virions in the 5’ UTR and observed similar patterns as those obtained in infected cells (in vivo) and protein-free solution (in vitro), confirming the RNA stability in this region (54).

**Figure 7.**
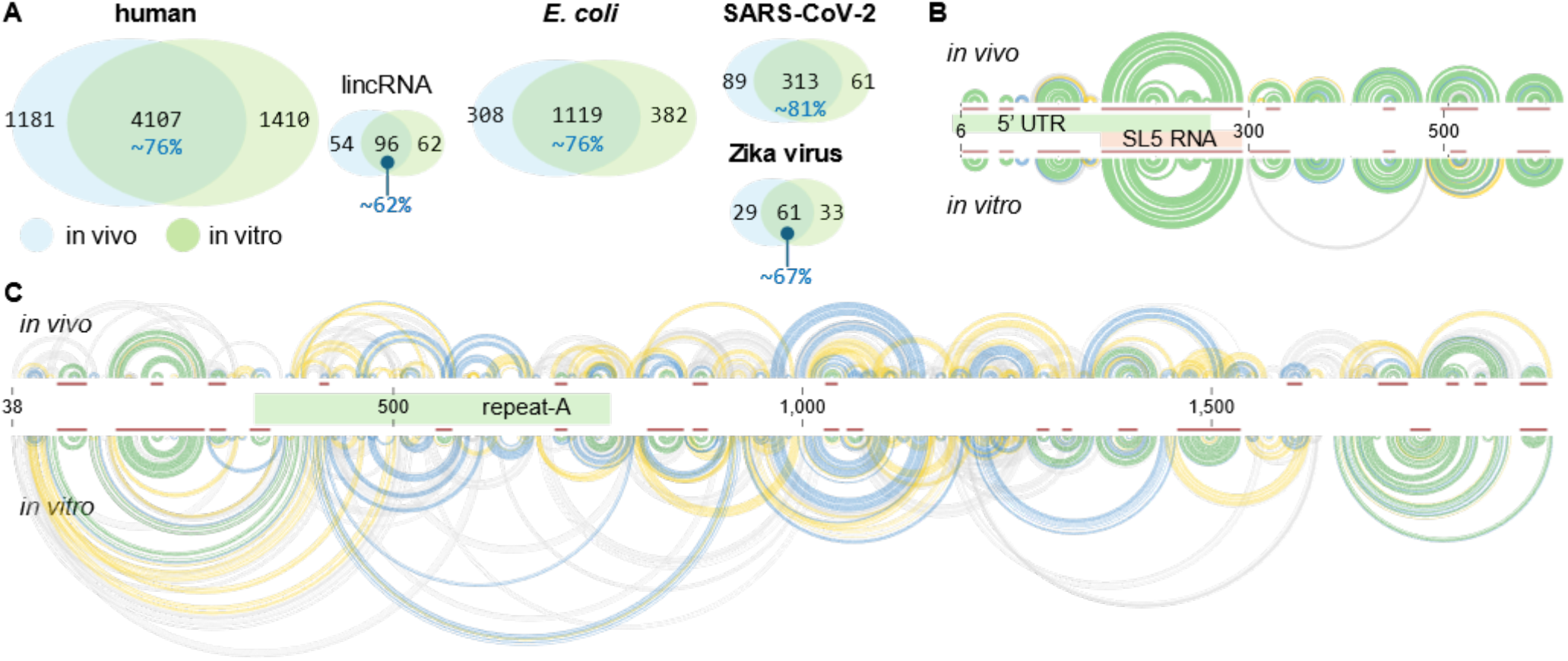
**(A)** Venn plots of overlapped stable sequences in the in vivo and in vitro data sets for humans, *E. coli*, and viruses. **(B-C)** Arc plots representing the base-pairing within (B) the first ∼600 nts in the SARS-CoV-2 genome and (C) ∼1950 nts in the Xist transcript (Ensembl ID: ENST00000429829). The colour of the arc represents the base-pairing probabilities: green (80–100%), blue (30–80%), yellow (10–30%), and grey (<10%). The identified stable sequences are highlighted in red bars in the middle of the plots.

In humans, the only annotated RNA type that has a smaller percentage of shared structures between in vivo and in vitro datasets (<70%) is long intergenic non-coding RNAs (lincRNAs) (Figure 7A). This is expected because lincRNAs are often associated with chromatin-modifying complexes (60), which are absent in the in vitro setting. In a prototype lincRNA Xist, the base-pairing patterns for in vivo and in vitro are distinct. For example, in the first two thousand nucleotides at the 5’-end, dynamic RNA base pairing was observed; fewer than half of the stable structures identified are shared between the in vivo and in vitro datasets (Figure 7C). I observed particularly dynamic base pairing in the repeat-A region, a well-studied, GC-rich, conserved sequence in Xist. These results are consistent with previously reported small-angle X-ray scattering analysis (61) and chemical probing results (62). Interestingly, a recent study identified a small molecule that can bind to repeat-A both in vivo and in vitro via an inducedfit mechanism, indicating that the lack of stable RNA structures may not impede its druggability, although a more detailed study is needed.

### Three- and four-way junction RNA structures

In the R2S-Finder pipeline, the criteria I chose to identify common stable secondary structures (see Material and Methods) precluded the identification of most complex tertiary or high-order structures, such as pseudoknots. However, my approach is capable of identifying three- and four-way junctions, where two and three loops exist in a single stable region, respectively. I determined that for all organisms, <30% of three- and four-way junctions are shared between in vivo and in vitro datasets. The low percentage of shared RNA junctions between in vivo and in vivo data suggests that RNA-binding proteins or other cellular factors influence the formation of these complex structures in vivo. The shared junctions that do appear in both conditions are likely more intrinsically stable, driven primarily by RNA folding rather than external factors, making them attractive candidates for drug targeting. I selected some representative three- and four-way junctions and predicted the 3-dimensional (3D) structures using transform-restrained Rosetta for RNA (trRosettaRNA), a computational tool used to predict RNA 3D structures by integrating evolutionary information and distance restraints (40) (Figure 8). I further predicted potential pockets for each structure using the Fpocket package, which identifies potential ligand-binding locations within these structures. (41). Some of the predicted pocket spaces were located in the central junction region, making them potential unique targets for RNA-targeting drug development. One of the stable four-way junction structure identified, SL5 in SARS-CoV-2’s 5’ UTR, includes a single G bulge that my laboratory previously identified as the binding site for a series of ligands (e.g., C30) (17). The proximity of the potential binding sites compared to the G bulge opens the opportunity to use these ligands in conjunction to improve RNA selectivity. For example, bivalent approaches simultaneous target two different binding sites within an RNA structure, which could enhance the selectivity and efficacy of the therapeutic compound by ensuring stronger and more specific binding.

**Figure 8.**
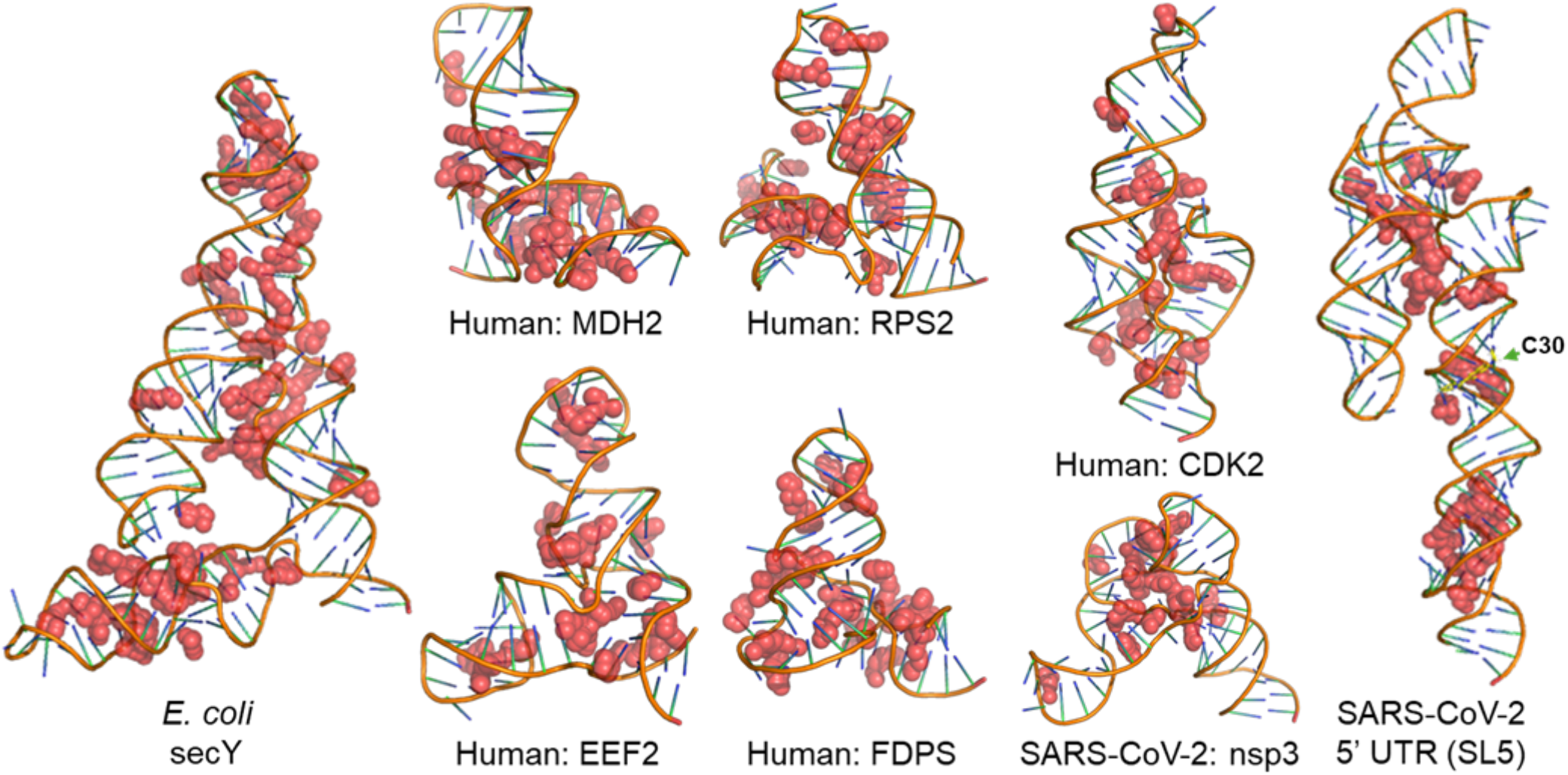
Representative 3D structures (predicted by trRosettaRNA, except for SARS-CoV-2’s SL5 RNA) of the stable three- and four-way junctions in different organisms. The potential binding pockets of these structures (predicted by Fpocket) are highlighted in red balls. The SARS-CoV-2 5’ UTR binder C30 (yellow) was aligned with SL5 RNA (Model Archive: Q6HL4). SL5 RNA structure was predicted in a previous computation report (63).

## DISCUSSION

In this study, I utilized published genome-wide chemical probing data available in RASP to provide a visualizable atlas of stable RNA secondary structures across multiple organisms. Given that RNA secondary structures play a crucial role in regulating RNA processes and can be targeted by synthetic small molecules, the information provided here is valuable for understanding the selectivity of such drug development endeavors.

This study has several limitations. Firstly, the coverage of SHAPE scores in the published datasets is limited. In humans, data from only approximately 1,300 genes were extracted, which represents only about 5% of the whole genome. In *E. coli*, the coverage is also around 5%. Due to this limited coverage, the observations in this study may be biased towards SHAPE(+) sequences, where the RNA is more accessible to chemical probing. Additionally, this lack of comprehensive coverage means the database may not be suitable for directly predicting off-target effects of RNA-binding drugs. Although the coverage of SHAPE scores is relatively low, the statistics of stable sequences, stem-loops, internal loops, and bulges identified remain highly informative; for example, we identified the prevalence of AU-rich tetraloops in both humans and *E. coli*. We also observed that 5’ UTRs in humans and viruses have more stable sequence ratios compared to the CDS regions and 3’ UTRs.

In the past decade, powerful algorithms have been developed to use chemical probing data to predict RNA conformations in equilibrium, such as RNA interaction groups by mutational profiling (RING-MaP) (64) and detection of RNA folding ensembles using expectation-maximization (DREEM) (65). It is possible that a more comprehensive dataset could be generated for dynamic structures of equilibria of multiple conformations, but this would require significantly more computational power. Despite these shortcomings, this study provides novel insight into the prevalence of basic RNA structural elements across organisms using genome-wide chemical probing data and provides a resource for further investigations. This dataset is especially useful for estimating the extent of off-target effects in humans for chemicals that target RNA secondary structures, and it could guide the development of more selective RNA-targeting drugs by identifying less commonly occurring RNA structures. Future work could focus on expanding this dataset to include more organisms and further refining our understanding of RNA structural dynamics, which could open up new avenues for therapeutic RNA targeting.

## CONCLUSION

This study presents a systematic approach for identifying and analysing stable RNA secondary structures across multiple organisms using genome-wide chemical probing data. By integrating chemical probing data with RNA folding models, I successfully identified stable sequences, including stem-loops, internal loops, bulges, and three- and four-way junctions in the genomes of humans, *E. coli*, SARS-CoV-2, and Zika virus. The analysis highlighted the prevalence of key structural motifs, such as tetraloops and 1×0 bulges, which may play critical roles in RNA stability and function. Comparisons between in vivo and in vitro conditions revealed notable differences in RNA folding, particularly in non-coding RNAs and viral genomes, emphasizing the influence of cellular environments on RNA structure. Furthermore, the discovery of potential drug-binding pockets within the identified stable RNA junctions, including the SARS-CoV-2 SL5 region, offers promising avenues for the development of RNA-targeting therapeutics. Overall, this study not only enhances our understanding of RNA structural dynamics but also provides a valuable foundation for future research on RNA-based drug discovery.

## SUPPLEMENTARY DATA

Codes for stable structure determination and full data sets of identified loops and bulges are available on the GitHub page: https://github.com/JingxinWangLab/R2S-Finder.

## ACKNOWLEDGEMENT

I thank Sarah Sumner at the University of Chicago for writing assistance.

